# A metagenomic approach to the molecular analysis of bacterial communities in tailings of a gold mine

**DOI:** 10.1101/2022.06.01.494447

**Authors:** Somayeh Parsania, Parisa Mohammadi, Mohammad Reza Soudi, Sara Gharavi

**Author notes:** Corresponding author. **E-mail:** (PM).

## Abstract

Gold mine operations release arsenic pollutants in environment. The present study investigated the diversity of bacterial communities in the arsenic-contaminated tailings dam effluent (TDE) of Zarshuran gold mine, Takab, northwest of Iran. The bacterial communities were examined using the next-generation sequencing method (Illumina) targeting the V3-V4 region of 16S rRNA genes. *Bacteroidetes* (50.3%), *Proteobacteria* (45.49%), *Actinobacteria* (1.14%) and *Firmicutes* (1.08%) constituted dominant phyla in the TDE. Its diversity was analyzed, and compared with that of three previously-studied arsenic-contaminated groundwater (GW) microbiomes. The raw sequencing data were analyzed in QIIME2. The prevalent taxonomic groups observed in all of the samples belonged to *Proteobacteria* (8.06-45.49%), *Bacteroidetes* (1.85-50.32%), *Firmicutes* (1.00-6.2%), *Actinobacteria* (0.86-5.09%), *Planctomycetes* (0.05-9.37%) and *Cyanobacteria* (0.6-2.71%). According to Principal Coordinates Analysis (PCoA), the TDE and GW samples were divided into high and low groups, based on their arsenic content, respectively. The average numbers of observed alpha diversity indices, i.e. Pielou’s evenness and Faith’s phylogenetic diversity, were not significantly different (P=0.18, Kruskal-Wallis test). According to the beta diversity analysis (unweighted), the observed operational taxonomic units (OTUs) and the diversity of the bacterial populations were not significantly different in the TDE, compared to the GW samples (PERMANOVA, P=0.276, 999 Permutations).

## 1. Introduction

Arsenic (As) constitutes the most prevalent environmental toxin in aquatic and terrestrial environments. This pollution primarily forms as inorganic oxyanions, arsenate (As^5+^) and arsenite (As^3+^). As^5+^ is the dominant species in oxygenated surface water, whereas As^3+^ is observed in anoxic or reduced conditions and is more toxic and soluble than As^5+^ [1]. Arsenic in ecosystems naturally originates from the environment or anthropogenic sources [2, 3]. Mining is the main source of arsenic pollutants in the environment [4]. Human industrial activities and natural processes such as rock weathering discharge arsenic into the surrounding environment. Arsenic in water and soil can enter the biogeochemical cycle and food chain, accumulate in the human body and pose different risks to human health [5].

Different mechanisms used by bacteria to neutralize arsenic toxicity include oxidation of As^3+^, reduction of As^5+^ and methylation and demethylation of arsenical compounds [6-8]. Bacteria can change the chemical properties, bioavailability, toxicity, natural behavior and environmental kismet of arsenic [6]. The expression of genes involved in arsenic metabolism is highly regulated by bacteria and controlled by operons such as *aio, ars* and arr [9, 10]. Arsenic-resistant bacteria utilize multiple arsenic-resistance operons for arsenic detoxification in complex and polluted environments [11].

Ecological research on mining waste suggests the richness and diversity of microorganisms are significantly changed in these areas. Numerous studies showed the adverse effects of metalloids on microbial communities in the soil. The short-term effects of heavy metals include decreases in the microbial community. Heavy metals also inhibit the growth of metal-sensitive microbial biomass. Moreover, prolonged exposure to heavy metals causes natural selection and expression of the most compatible genes with existing conditions [12].

Culture-independent molecular methods concerning 16S rRNA genes can characterize microbial ecology [13]. High-throughput sequencing such as Illumina can also be used as a promising and rapid method to analyze gold mines’ microbial diversity and community structure [14-17]. The present study employed the high throughput sequencing technology to investigate the diversity of the bacterial community in the arsenic-polluted TDE of Zarshuran gold mine [18]. The diversity was analyzed in three arsenic-contaminated groundwater (GW) microbiomes. The differences in the richness and diversity of bacterial communities were assessed between the TDE and GW samples. Major microbial activities of the arsenic cycle in the TDE were characterized.

## 2. Materials and methods

### 2.1. Site description and sample collection

Zarshuran gold mine (36°41’02.7”N, 47°08’05.6”E) is located in West Azerbaijan Province, Northwestern Iran. The main types of minerals in this area are orpiment (As_2_S_3)_ (> 60%), realgar (As_2_S_4_) and arsenopyrite (FeAsS) [19], which are soluble minerals in alkaline solutions. Different methods such as cyanidation are used for gold mining. This leads to the dissolution of orpiment mineral ores in alkaline solutions and releases arsenic species in tailings dam effluents [19]. The arsenic-contaminated wastewater was collected from TDE of Zarshuran gold mine in June 2018.

For metagenomics analysis, 5 liters of the TDE were collected in sterilized polypropylene bottles and straightly transferred to the laboratory. For physicochemical and heavy metals analysis, sampling was performed according to the standard methods for examining water and wastewater [20]. Moreover, heavy metal contents were assayed by the polarography method (Metrohm-Model 797 VA). Polarography analysis was performed for arsenic elements in the range of –0.7 to –0.55 V. Simultaneous detection of Ni-Co and Zn-Cu-Pb-Cd was conducted in the range of -1.25 to -0.8 V and -1.25 to +0.05 V, respectively, using dropping mercury electrodes. Solutions were de-oxygenated with a stream of nitrogen gas (1atm).

### 2.2. Genomic DNA extraction and DNA amplification and sequencing

Five litres of TDE sample were filtered through 0.22μ sterilized nitrocellulose filters (Millipore), and the filters were stored at -80 °C in a deep freezer until further analysis. According to the manufacturer’s protocol, the DNA of the collected samples was extracted using DNeasy PowerWater Kit (Qiagen, Germany). The DNA samples were stored at -20 °C until further processed. Universal bacterial primers 27F and 1492R were used to amplify the full-length 16S rDNA for environmental microbiome surveys. DNA concentration and purity were analyzed by agarose and NanoDrop 2000 (Thermo, USA). Approximately 80 ng/μl of purified PCR product of sample was sent to Macrogen, Inc (Seoul, South Korea) for a second PCR amplification and sequencing. The second PCR amplification of the V3-V4 region of the 16S rRNA genes was carried out using the universal primers, 341F/785R. The PCR amplicons were sequenced by paired-end sequencing chemistry on the Illumina MiSeq, Macrogen, Inc. The raw sequencing data file was conceded to the NCBI Sequence Read Archive (SRA) (accession number: PRJNA721938).

### 2.3. Analysis of community by 16S rRNA gene

Read data was assayed using QIIME 2 (https://qiime2.org), as provided online manuals (https://docs.qiime2.org/2019.10/tutorials/moving-pictures/) [21]. Diversity composition analysis of TDE was assessed with three arsenic-contaminated GW. Microbiome (G3, G6 and G19) from Rayong in Thailand, in which the bacterial diversity centered on the V3-V4 region of the 16S rRNA gene of which was previously studied [22] and their FASTQ files, (G3 (SRR6760375), G6 (SRR6760374) and G19 (SRR6760381)) were available in ENA database. The sequence quality control and table construction of feature were made using DADA2 (–p-trunc-len-f 250–p-trunc-len-r 240–p-trunc-q 15–p-trim-lef-f 11–p-trim-lef-r 16) [23]. The settings for quality control were calibrated according to the reads’ quality distribution over the length of the sequence. These sequences were grouped into the operational taxonomic units (OTUs) based on a 100% similarity threshold. Alpha rarefaction analysis, the taxonomic classification of OTUs, alpha diversity (the number of observed OTUs, Shannon diversity, and Faith’s phylogenetic diversity) and beta diversity (Jaccard distance, Bray-Curtis distance, unweighted and weighted UniFrac distances) were analyzed using QIIME 2. For taxonomic classification, Greengenes 99% OTUs (13_8 release) were utilized as 16S rRNA gene databases [24]. The taxonomic classifier was carried out in QIIME 2, originally trained by 341F/785R region of Greengenes registered sequences. The statistical assessment for diversity metrics and creation of principal coordinate analysis (PCoA) schema were calculated. Furthermore, beta diversity metrics were performed through QIIME 2. Sampling depth was set to 98515. The heat map is generated concerning on Qiime taxa filter-table and Qiime feature-table to retain only features which were annotated to the genus level by the most frequent (>300 reads) in datasets.

## 3. Results

### 3.1. TDE geochemistry

Arsenic levels, pH, Total Organic Carbon (TOC), Electrical Conductivity (EC). and heavy metals such as Zn, Cu, Pb and Ni were measured in the TDE. The physicochemical and heavy metal analyses of the TDE sample showed significant concentrations (μg/l) of zinc (71.5), copper (108.2), lead (59.7), nickel (20.1), cobalt (4.9), cadmium (<0.471) and arsenic (7750). The physicochemical parameters were obtained as pH:8.02, conductivity:56.98 μS/cm and TDS:36.98 mg/l. Parameters associated with physicochemical properties and heavy metals in the GW samples were adopted from literature (Table 1) [22].

**Table 1.**
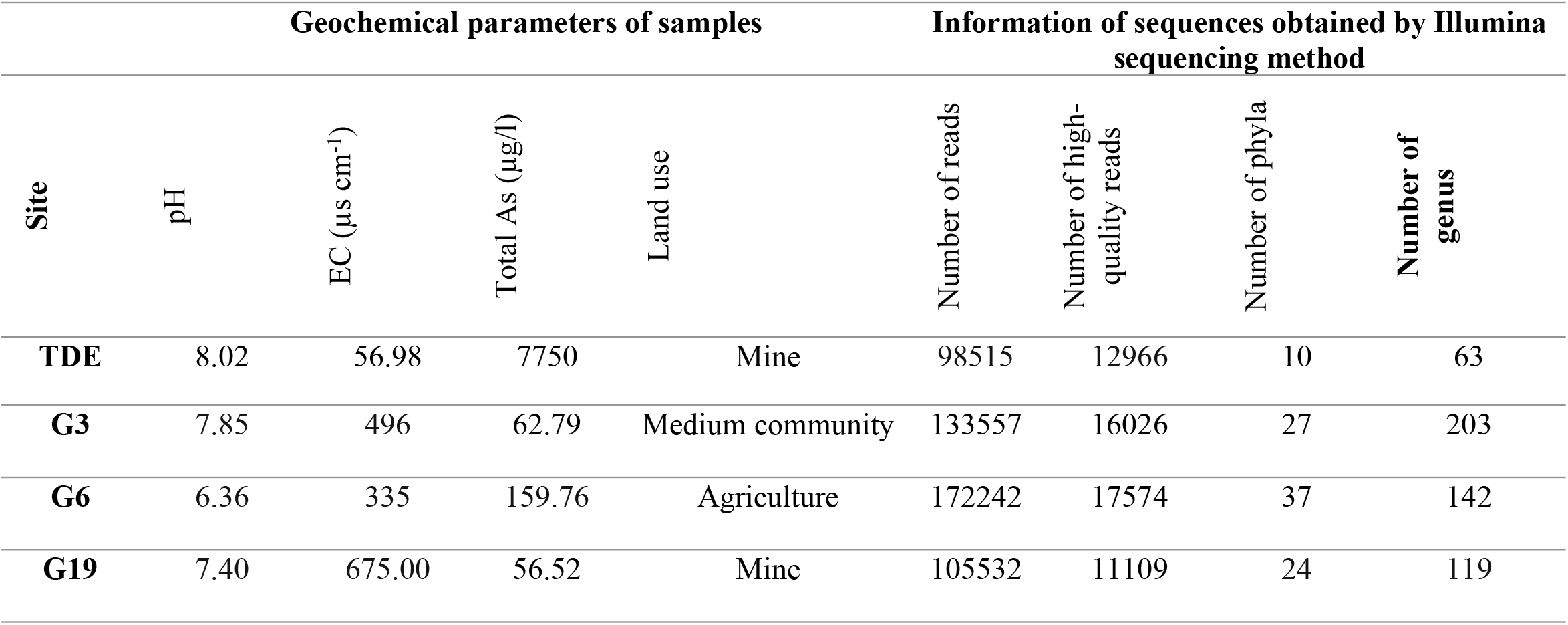
The geochemical parameters and sequence information of the samples

### 3.2. Richness and evenness

The Illumina-MiSeq analysis of the V3-V4 region of bacterial 16S rRNA genes showed a mean Phred quality score of 30 (Phred Q30) in 82.38% of 197,030 paired-end reads. After removing primers and chimeras with an average read length of 250 bp and filtering the length/quality (quality threshold>15), 11109-17547 reads were obtained with a mean frequency of 14418.75. A plateau was observed at a sequencing depth 98515 on the Shannon-Wiener curve of all the samples, suggesting the depth adequacy of sequencing (Fig 1). An increase in the sequence number was also observed on the rarefaction curve of each sample as a function of the OTUs. The plateau observed on the rarefaction curve showed that a significant portion of diversity was captured through metagenomic efforts.

**Fig 1.**
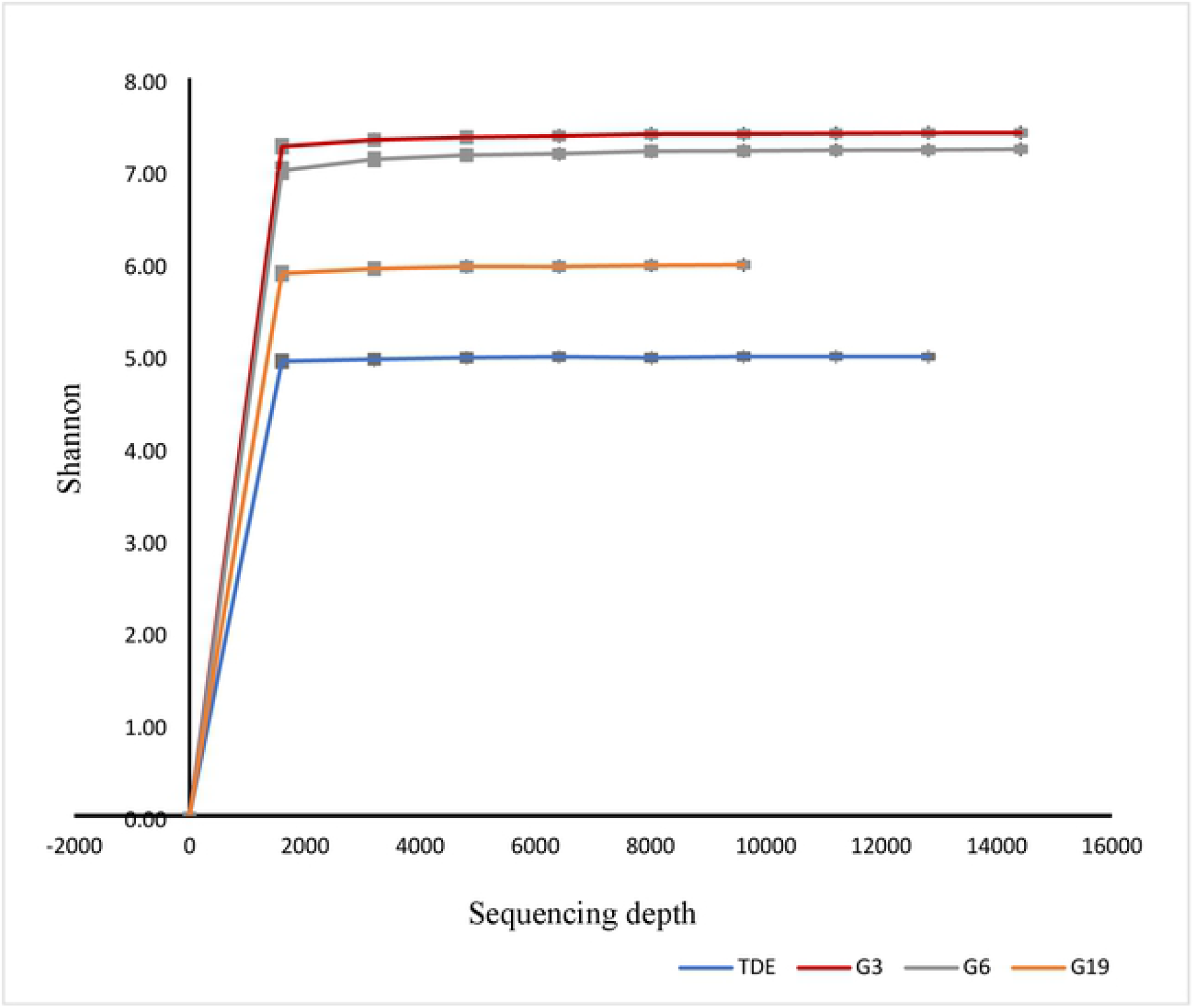
Shannon-Wiener curves of each sample; these curves were all calculated at 99% level of similarity with Illumina Miseq data for microbiomes of TDE and GW samples.

### 3.3. Bacterial diversity

The QIIME2 analysis showed 1191 OTUs in four samples, suggesting that 99.94-100% of the reads were assigned to bacteria and 0-0.06 % to Archaea. Forty-five bacterial phyla and 441 bacterial genera were retrieved from the samples. Archaeal sequences were affiliated to *Euryarchaeota, Parvarchaeota* and *Crenarchaeota* and only detected in GW samples. Common sequences of the prevalent taxonomic groups in all the samples were affiliated to *Proteobacteria* (8.06-45.49%), *Bacteroidetes* (1.85-50.32%), *Firmicutes* (1.00-6.2%), *Actinobacteria* (0.86-5.09%), *Planctomycetes* (0.05-9.37%) and *Cyanobacteria* (0.6-2.71%) (Fig 2). As the main phylum (3 out of 4), *Proteobacteria* accounted for 62.21% of total valid features.

**Fig 2.**
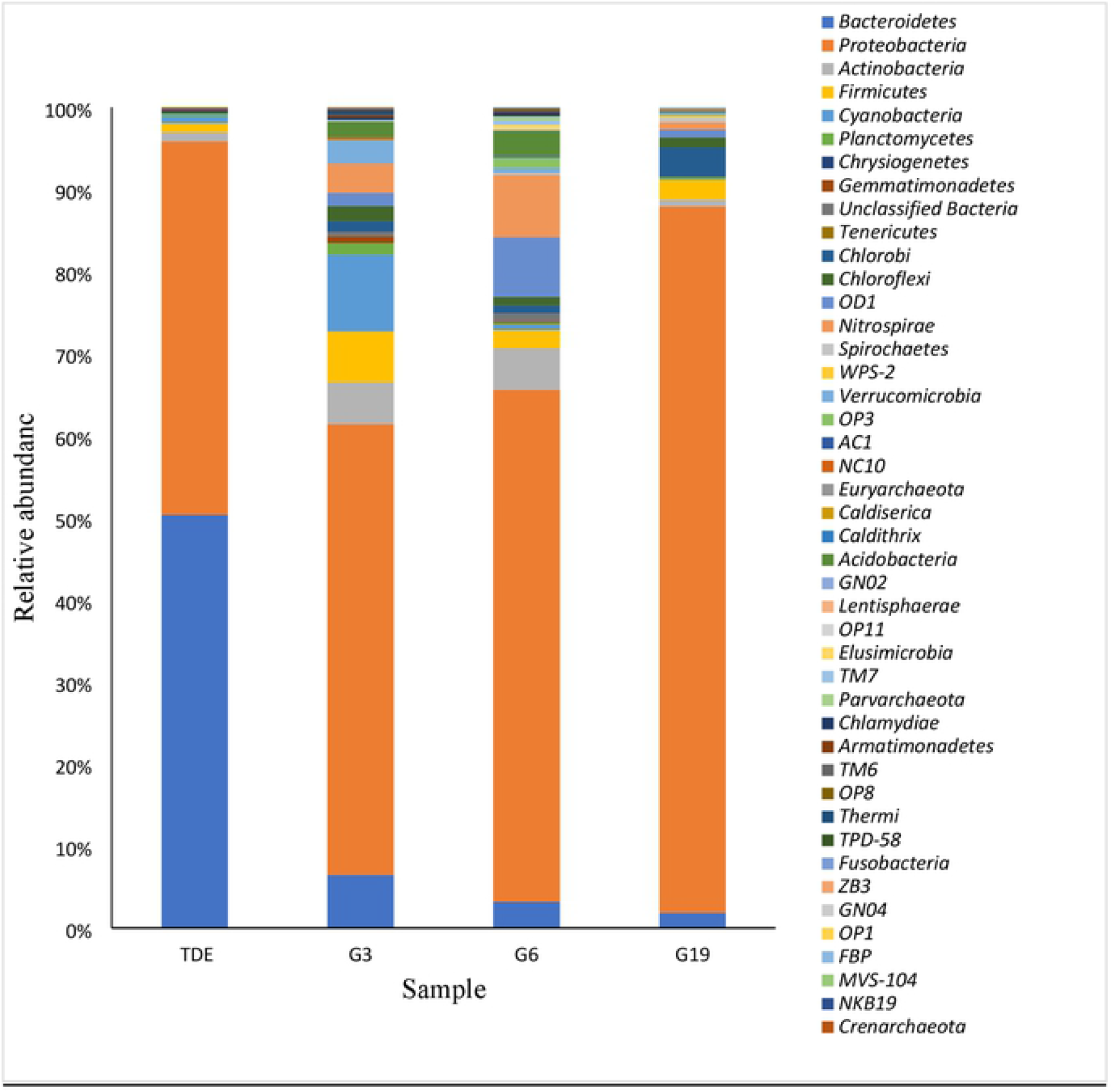
The taxonomic compositions at the level of phylum

The TDE primarily comprised *Bacteroidetes* (50.3%), *Proteobacteria* (45.49%), *Actinobacteria* (1.14%) and *Firmicutes* (1.08%), which was inconsistent with the G3 sample containing phyla *Proteobacteria* (54.93%), *Cyanobacteria* (9.37%), *Bacteroidetes* (6.46%) and *Firmicutes* (6.2%). A shift was observed in the G6 sample and four abundant phyla comprised *Proteobacteria* (62.38%), *Nitrospirae* (7.6%), *OD1* (7.3%) and *Actinobacteria* (5.08%). *Proteobacteria* (86.06%) followed by *Chlorobi* (3.57%), *Firmicutes* (2.3%) and *Bacteroidetes* (1.85%) were the most abundant phyla in the G19 sample collected from areas surrounding the mine.

A total of 441 genera obtained from all the samples included 203 from G3, 142 from G6, 119 from G19 and 63 from the TDE. Table 2 presents dominant bacterial genera among the four samples (≥1% of total OTUs). Richness slightly decreased at the genus level in G3, G6, G19 and TDE samples. Six dominant genera were retrieved from the TDE sample, 178 from G3, 8 from G6 and 6 from G19. Members of *Erythrobacteraceae* (10%), *Rhodobacteraceae* (1%) and *Beijerinckiaceae* (1.08%) and order of *Sphingobacteriales* (1.02%) observed in the TDE sample were not identified at genus or family levels, suggesting unknown or unclassified bacteria in the TDE samples.

**Table 2.**
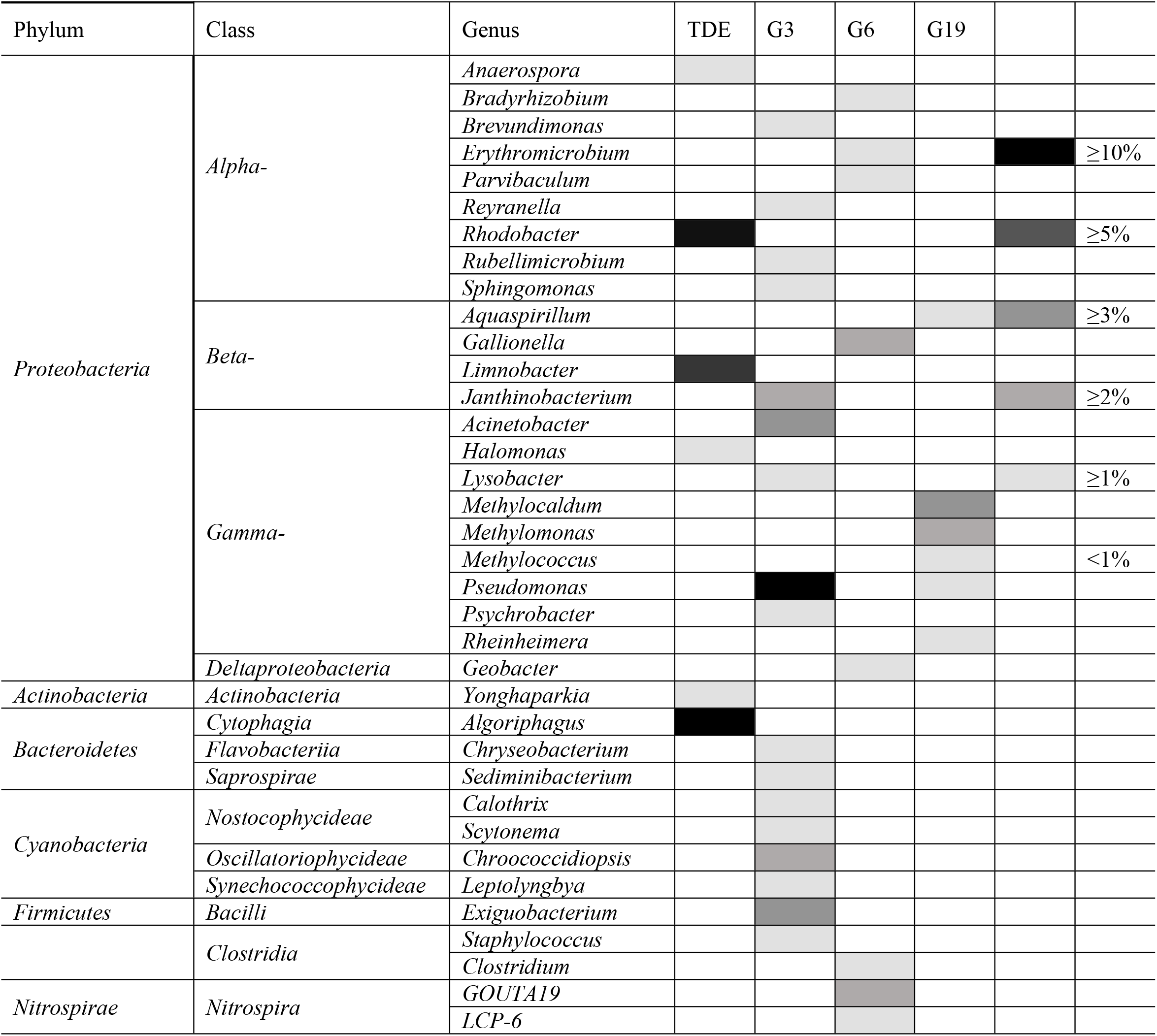
The bacterial genera were retrieved in the arsenic-contaminated TDE and GW samples.

A higher abundance of over 1% of the total sequences in the TDE sample was observed only in six genera, including *Algoriphagus* (*Bacteroidetes*), *Rhodobacter* and *Anaerospora* (*Alphaproteobacteria), Limnobacter* (*Betaproteobacteria), Halomonas* (*Gammaproteobacteria)*, as well as *Yonghaparkia (Actinobacteria)*, as a unique dominant genus in the TDE sample. The most abundant genera included *Algoriphagus (*45%), *Rhodobacter* (12.7%) and *Limnobacter* (5.3%).

The present research found significant differences between the TDE and GW samples in terms of their microbial community structure. *Pseudomonas* and *Acinetobacter* in G3 and *Acinetobacter* in G19 were highly abundant in the low-arsenic GW sites. The vast majority of reads in G6 were related to an unidentified genus. The rarely-observed bacteria included *Methylosinus* in all the samples, *Pseudomonas* in TDE, G3 and G19 and *Bacillus* and *Nocardioides* in TDE, G3 and G6. The samples were, however, different in terms of their abundant genus. Fig 3 shows the heat maps of the main feature (>300 reads) in datasets.

**Fig 3.**
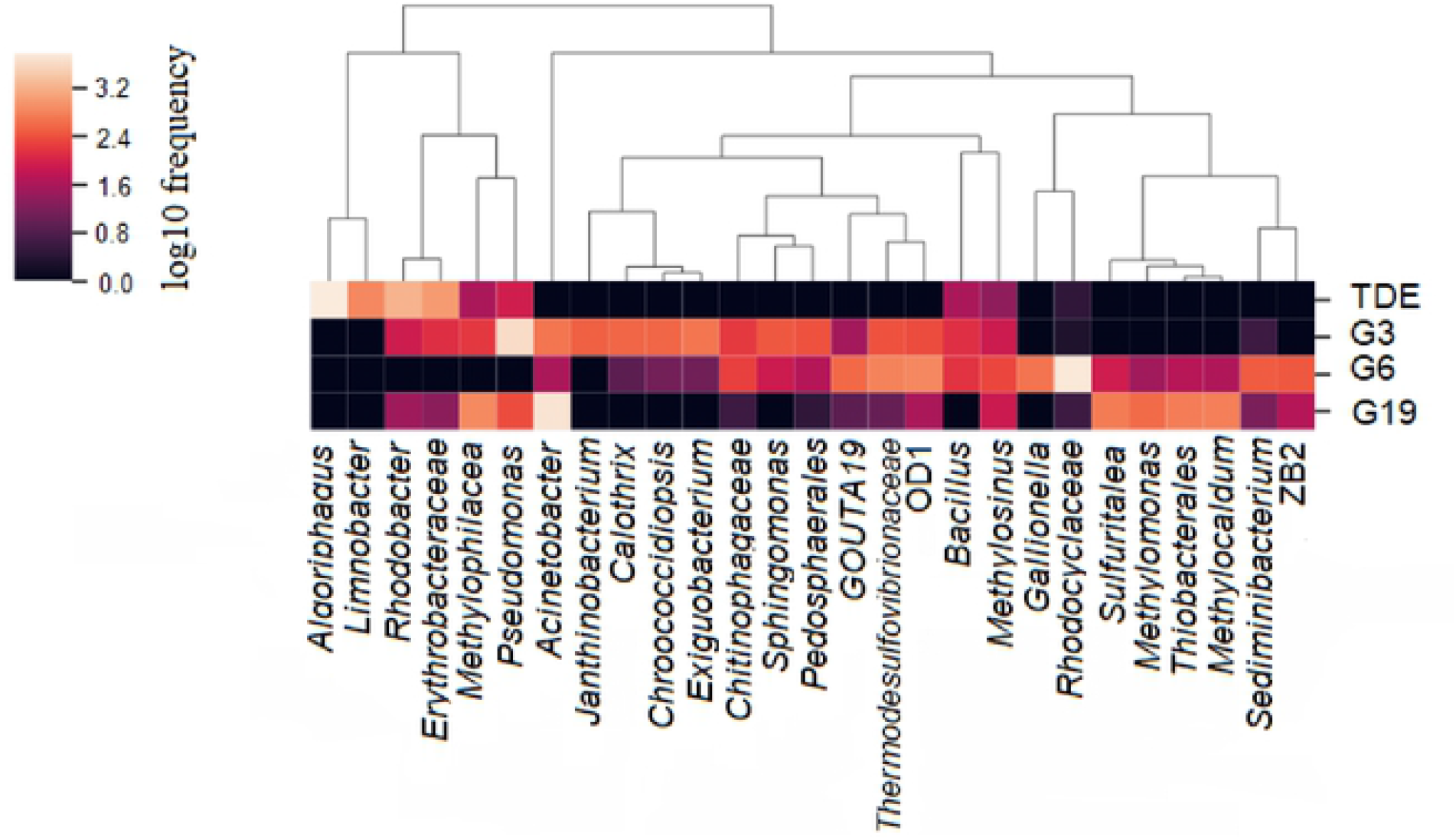
Heat map of the most frequent feature (>300 reads) in datasets (TDE, G3, G6 and G19). Darker or lighter color represent lower or higher frequency, respectively.

### 3.4. Bacterial Community Structure

The TDE and three GW samples were divided into two groups according to PCoA. The first group including the TDE sample was characterized by a high concentration of arsenic (7.75 mg/l). The other group included three GW samples (G3, G6 and G19) with low arsenic concentrations (62.79 μg/l, 159.76 μg/l and 56.52 μg/l, respectively) (Table 1). The average numbers of the observed alpha diversity indices of Pielou’s evenness and Faith’s phylogenetic diversity were not significantly different (P=0.18, Kruskal-Wallis test). Pielou’s evenness and Faith’s phylogenetic diversity measured the evenness and richness of the community, respectively. The beta diversity analysis (unweighted) also showed insignificant differences between the TDE and GW samples in terms of OTUs and diversity of bacterial population (PERMANOVA, P= 0.276, 999 permutations). Fig 4 shows principal coordinates of Jaccard and Bray-Curtis distance as the qualitative and quantitative measures of community dissimilarity.

**Fig 4.**
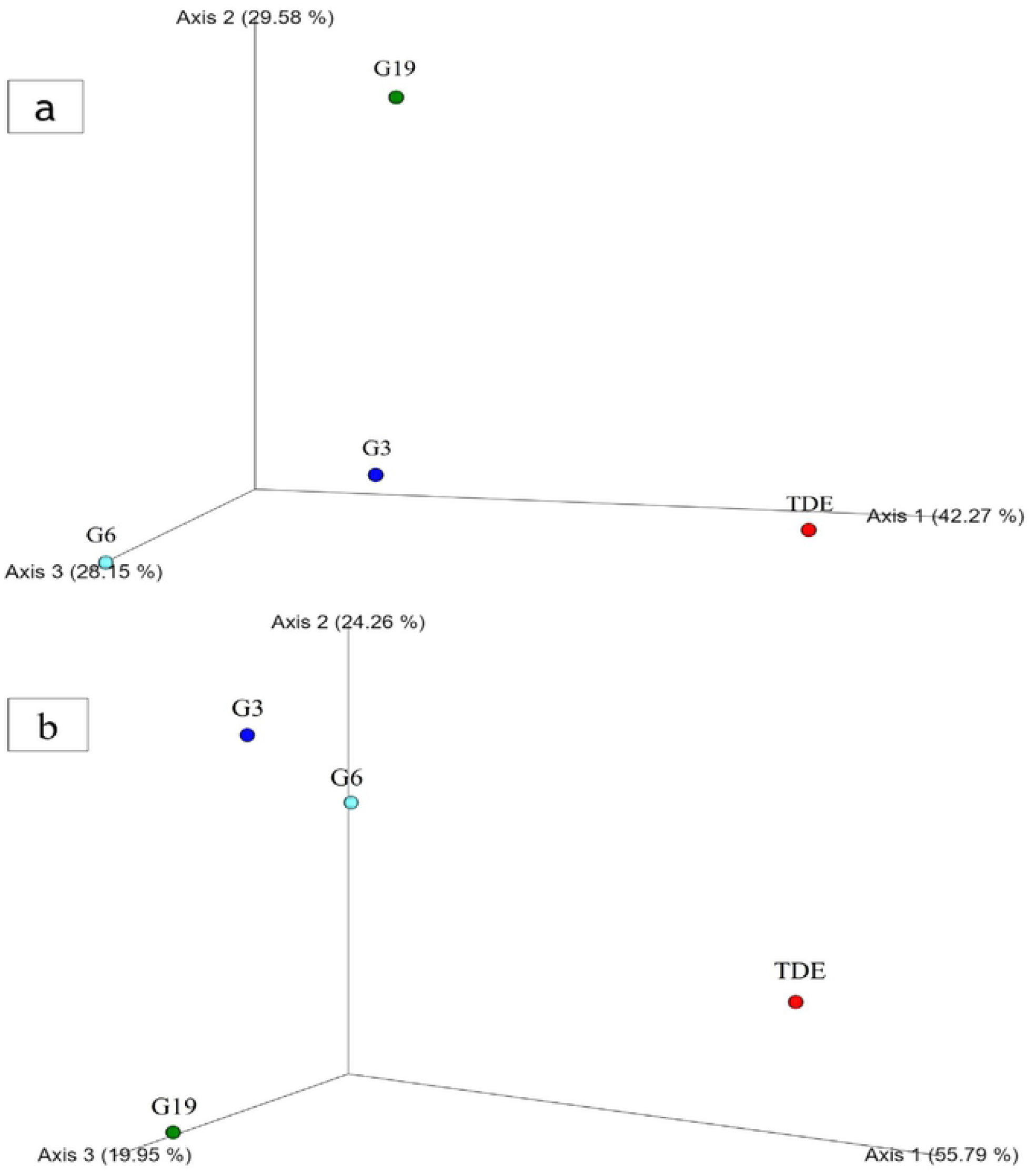
PCoA analysis of arsenic polluted water, (a) PCoA plots of unweighted-unifrac. Each point indicates to an individual sample and clustering of the points means the similarity in the components of OTUs among those samples. (b) PCoA plots of weighted-unifrac.

## 4. Discussion

Mine tailings contain limited organic substances [25], and microbiological activities are influenced by the type and chemistry of metal(loid)s [26]. The present results confirmed previous findings suggesting *Proteobacteria, Bacteroidetes, Firmicutes, Actinobacteria, Chloroflexi, Cyanobacteria, Acidobacteria, Gemmatimonadetes* and *Planctomycetes* constitute the main phyla in gold mine tailings contaminated with arsenic oxyanions [16, 27].

The TDE was less abundant in diversity (10 groups) at the phylum level than the GW samples, which can be explained by the high concentrations of arsenic, cyanide, antimonite and other toxic heavy metals and the limited number of available electron donors and acceptors in the TDE [27]. This low bacterial diversity in the TDE is consistent with previously-reported findings [28-30]. A low bacterial diversity was observed in the acidic drainage of the mine, which contained arsenic concentrations as high as 750-2700 mg/l.

The feature analysis of the NGS library showed *Proteobacteria* with a high frequency in all the samples irrespective of the arsenic components, which can be explained by the resistance or the tolerance mechanisms of *Proteobacteria* [12]. Numerous studies found *Proteobacteria* to be the main phylum [31-33]. The present study found *Bacteroidetes* to be the main phylum in the TDE. Bacterial communities isolated by Riley et al. from a river in South Africa contained different anthropogenic contaminants, including arsenic, chromium, nickel and uranium. The phylum-level concentrations of contaminants were significantly lower in downstream compared to in upstream sites. *Firmicutes, Acidobacteria, Cyanobacteria* and *Verrucomicrobia* were prominent at downstream sites, while *Bacteroidetes* was more frequently observed in the contaminated areas [34]. The high frequency of *Bacteroidetes* can be attributed to the harsh environment of the TDE and resistance or tolerance of this group of bacteria. Investigating the susceptibility of 105 strains of *Bacteroidetes* to heavy metal ions (As, Ag, Ni, Co, Pb, Cd, Cr and Hg) found all strains of *Bacteroidetes* to be multiple resistant [34]. *Bacteroides vulgatus* ATCC 8482 is highly resistant to As^5+^ and methyl arsenate. The arsenical-inducible transcriptional unit (*ars* operon) in the genome of *B. vulgatus ATCC* 8482 includes eight continuous genes. Furthermore, *arsR* and *arsDABC* genes of this operon play a key role in the detoxification of arsenic [35]. About 0.2% of the sequence reads of the OTUs also belonged to the unclassified bacterial taxa, which showed unknown microorganisms in this TDE sample.

*Algoriphagus, Rhodobacter* and *Limnobacter* are the most abundant genera in the TDE sample. *Algoriphagus* identified in shallow aquifers [36] and sediments [37] with high arsenic levels using metagenomic methods can reduce As^5+^ to As^3+^ using nitrate, acetate, sulfide and Fe(II) as electron donors.

As facultative anaerobes and photosynthetic bacteria, purple nonsulfur bacteria (PNSB) are flexible in carbon source utilization and resist to As^5+^, As^3+^ and other heavy metal ions. Different resistance mechanisms used by many PNSB and several other bacteria include conversion of metals to a reduced toxic state through modifying their oxidation state, arsenate reduction, arsenite methylation pathways [11, 38, 39] and efflux of heavy metals out of the cells [40]. Under anaerobic conditions, *Rhodobacter* can respire A_+5_ and oxidize certain carbon sources for hydrogen photofermentation [41]. Certain *Rhodobacter* species can generate exopolysaccharides with arsenic and lead chelating activities [42]. The high abundance of *Rhodobacter* in the harsh conditions of the TDE showed the potential of this organism for both bioremediation and use of effluents contaminated with arsenic and other heavy metals as a substrate.

As a sulphur-oxidizing bacterium, *Limnobacter* was the third abundant genus in the TDE sample, suggesting a feasible practical biogeochemical sulphur cycle in this area [43]. The abundance of *Limnobacter* in the TDE of Zarshuran mine ores containing sulphide minerals was not unexpected. This genus comprises species such as *Limnobacter litoralis* [44] and *Limnobacter thiooxidans* [45] which can oxidize thiosulfate. Although this study did not investigate thiosulfate, *Limnobacter* spp. can play a role in sulfur or thiosulfate oxidation. *Anaerospora* is a genus with sulfur and/or iron-reducing potential. *Anaerospora hongkongensis* can perform ferrihydrite reduction and anaerobic oxidation of ammonium driven by sulfur redox cycling [46]. Observing this group of bacteria was not unexpected given the use of ammonia as an oxidizing agent for sulphur-bearing minerals in gold mining. This genus can be also involved in the arsenic cycle given that it was identified in paddy soil bacterial communities at different heavy metal levels [47]. The bacteria belonging to the *Yonghaparkia* genus and isolated from microbial communities in a uranium mine (Athabasca Basin, Canada) [48], a gold mine (Linglong, China) [15] and the black shale (China) [49] can contribute to the biogeochemical cycles of S and Fe.

As an arsenic-resistant gammaproteobacterium, *Halomonas* sp. was frequently observed in both the TDE sample and arsenic-contaminated environments. *Halomonas* sp. strain GFAJ-1 was found to use arsenic as an alternative to phosphorus to survive under phosphate deficiency conditions [50]. The arsenate reductase and arsenite oxidizing activities of *Halomonas* sp. play key roles in arsenic biogeochemical cycles [51]. *Halomonas* species contain arxA gene and can contribute to arx-dependent arsenite oxidation or couple As^3+^ oxidation with nitrate reduction [52]. The increased exopolysaccharide production observed in *Halomonas* sp. isolated from the rhizosphere under arsenic stress plays a key role in arsenic sequestration [53]. In line with literature, the present study found *Pseudomonas* and *Acinetobacter* isolated from aquifers with high arsenic levels to be involved in the arsenic cycle, including arsenic resistance, As^5+^ reduction and As^3+^ oxidation [54, 55].

## 5. Conclusion

This study was conducted on the bacterial diversity and community structure of a gold tailings dam. Metagenomic analyses showed *Algoriphagus, Rhodobacter, Anaerospora, Limnobacter, Halomonas* and *Yonghaparkia* to be the main bacterial genera. Despite the limited similarities in the prokaryotic community of the samples, most of the retrieved genera in the TDE are unique and we retrieved the native bacterial genus of Iran. The present study suggested the key role of functional bacteria in the geochemical cycles of heavy metals, which helps with the scientific management of the bioremediation and ecological risk assessments of gold mine tailings.

## Acknowledgements

The authors appreciate the Vice-chancellor of Alzahra University and the technicians of Shayesteh Sepehr Laboratories of Industrial Microbiology for their cooperation. We would like to especially thank Dr. Sarikhani for his scientific comments and discussion on the interpretation of NGS data.

## References

1. Shafiquzzaman M, Azam MS, Nakajima J, Bari QH. Investigation of arsenic removal performance by a simple iron removal ceramic filter in rural households of Bangladesh. Desalination. 2011;265(1-3):60–6. https://doi.org/10.1016/j.desal.2010.07.031

2. Al-Abed SR, Jegadeesan G, Purandare J, Allen D. Arsenic release from iron rich mineral processing waste: influence of pH and redox potential. Chemosphere. 2007;66(4):775–82. https://doi.org/10.1016/j.chemosphere.2006.07.045

3. Domingo JL. Prevention by chelating agents of metal-induced developmental toxicity. Reproductive Toxicology. 1995;9(2):105–13. https://doi.org/10.1016/0890-6238(94)00060-3

4. Bech J, Roca N, Tume P, Ramos-Miras J, Gil C, Boluda R. Screening for new accumulator plants in potential hazards elements polluted soil surrounding Peruvian mine tailings. Catena. 2016;136:66–73. https://doi.org/10.1016/j.catena.2015.07.009

5. Hasanuzzaman M, Nahar K, Fujita M. Mechanisms of Arsenic Toxicity and Tolerance in Plants: Springer; 2018. https://doi.org/10.1007/978-981-13-1292-2

6. Oremland RS, Stolz JF. The ecology of arsenic. Science. 2003;300(5621):939–44. Epub 2003/05/10. doi: 10.1126/science.1081903. PubMed PMID: 12738852. https://doi.org/10.1126/science.1081903

7. Zhang J, Cao T, Tang Z, Shen Q, Rosen BP, Zhao F-J. Arsenic methylation and volatilization by arsenite S-adenosylmethionine methyltransferase in Pseudomonas alcaligenes NBRC14159. Appl Environ Microbiol. 2015;81(8):2852–60. https://doi.org/10.1128/AEM.03804-14

8. Yang H-C, Rosen BP. New mechanisms of bacterial arsenic resistance. Biomedical journal. 2016;39(1):5–13. https://doi.org/10.1016/j.bj.2015.08.003

9. Suhadolnik ML, Salgado AP, Scholte LL, Bleicher L, Costa PS, Reis MP, et al. Novel arsenic-transforming bacteria and the diversity of their arsenic-related genes and enzymes arising from arsenic-polluted freshwater sediment. Sci Rep. 2017;7(1):1–17. https://doi.org/10.1038/s41598-017-11548-8

10. Cleiss-Arnold J, Koechler S, Proux C, Fardeau M-L, Dillies M-A, Coppee J-Y, et al. Temporal transcriptomic response during arsenic stress in Herminiimonas arsenicoxydans. BMC genomics. 2010;11(1):709. https://doi.org/10.1186/1471-2164-11-709

11. Zhao C, Zhang Y, Chan Z, Chen S, Yang S. Insights into arsenic multi-operons expression and resistance mechanisms in Rhodopseudomonas palustris CGA009. Frontiers in microbiology. 2015;6:986. https://doi.org/10.3389/fmicb.2015.00986

12. Das S, Jean J-S, Kar S, Liu C-C. Changes in bacterial community structure and abundance in agricultural soils under varying levels of arsenic contamination. Geomicrobiol J. 2013;30(7):635–44. https://doi.org/10.1080/01490451.2012.746407

13. Hirsch PR, Mauchline TH, Clark IM. Culture-independent molecular techniques for soil microbial ecology. Soil Biology and Biochemistry. 2010;42(6):878–87. https://doi.org/10.1016/j.soilbio.2010.02.019

14. Sibanda T, Selvarajan R, Msagati T, Venkatachalam S, Meddows-Taylor S. Defunct gold mine tailings are natural reservoir for unique bacterial communities revealed by high-throughput sequencing analysis. Science of The Total Environment. 2019;650:2199–209. https://doi.org/10.1016/j.scitotenv.2018.09.380

15. Li M, Tian H, Wang L, Duan J. Bacterial diversity in Linglong gold mine, China. Geomicrobiol J. 2017;34(3):267–73. https://doi.org/10.1080/01490451.2016.1186763

16. Das S, Bora SS, Yadav R, Barooah M. A metagenomic approach to decipher the indigenous microbial communities of arsenic contaminated groundwater of Assam. Genomics data. 2017;12:89–96. http://dx.doi.org/10.1016/j.gdata.2017.03.013

17. Ji H, Zhang Y, Bararunyeretse P, Li H. Characterization of microbial communities of soils from gold mine tailings and identification of mercury-resistant strain. Ecotoxicology and environmental safety. 2018;165:182–93. https://doi.org/10.1016/j.ecoenv.2018.09.011

18. Asadi H, Hale M. A predictive GIS model for mapping potential gold and base metal mineralization in Takab area, Iran. Computers & Geosciences. 2001;27(8):901–12. http://dx.doi.org/10.1016/S0098-3004(00)00130-8

19. Aazami M, Lapidus G, Azadeh A. The effect of solution parameters on the thiosulfate leaching of Zarshouran refractory gold ore. International Journal of Mineral Processing. 2014;131:43–50.

20. Rice E, Baird R, Eaton A, Odor T, By D, Carbon TO. Standard Methods for the Examination of Water and Wastewater. 2017. https://doi.org/10.1016/j.minpro.2014.08.001

21. Bolyen E, Rideout JR, Dillon MR, Bokulich NA, Abnet CC, Al-Ghalith GA, et al. Reproducible, interactive, scalable and extensible microbiome data science using QIIME 2. Nat Biotechnol. 2019;37(8):852–7. https://dx.doi.org/10.1038%2Fs41587-019-0209-9

22. Sonthiphand P, Ruangroengkulrith S, Mhuantong W, Charoensawan V, Chotpantarat S, Boonkaewwan S. Metagenomic insights into microbial diversity in a groundwater basin impacted by a variety of anthropogenic activities. Environmental Science and Pollution Research. 2019;26(26):26765–81. https://doi.org/10.1007/s11356-019-05905-5

23. Callahan BJ, McMurdie PJ, Rosen MJ, Han AW, Johnson AJA, Holmes SP. DADA2: high-resolution sample inference from Illumina amplicon data. Nature methods. 2016;13(7):581. https://doi.org/10.1038/nmeth.3869

24. DeSantis TZ, Hugenholtz P, Larsen N, Rojas M, Brodie EL, Keller K, et al. Greengenes, a chimera-checked 16S rRNA gene database and workbench compatible with ARB. Appl Environ Microbiol. 2006;72(7):5069–72. https://doi.org/10.1128/aem.03006-05

25. Gallego S, Esbrí JM, Campos JA, Peco JD, Martin-Laurent F, Higueras P. Microbial diversity and activity assessment in a 100-year-old lead mine. Journal of Hazardous Materials. 2021;410:124618. https://doi.org/10.1016/j.jhazmat.2020.124618

26. Dopson M, Johnson DB. Biodiversity, metabolism and applications of acidophilic sulfur-metabolizing microorganisms. Environ Microbiol. 2012;14(10):2620–31. https://doi.org/10.1111/j.1462-2920.2012.02749.x

27. Baker BJ, Banfield JF. Microbial communities in acid mine drainage. FEMS microbiology ecology. 2003;44(2):139–52. https://doi.org/10.1016/S0168-6496(03)00028-X

28. Bruneel O, Duran R, Casiot C, Elbaz-Poulichet F, Personné J-C. Diversity of microorganisms in Fe-As-rich acid mine drainage waters of Carnoules, France. Appl Environ Microbiol. 2006;72(1):551–6. https://doi.org/10.1128/AEM.72.1.551-556.2006

29. Delavat F, Lett M-C, Lièvremont D. Novel and unexpected bacterial diversity in an arsenic-rich ecosystem revealed by culture-dependent approaches. Biology direct. 2012;7(1):28. https://doi.org/10.1186/1745-6150-7-28

30. Guan X, Yan X, Li Y, Jiang B, Luo X, Chi X. Diversity and arsenic-tolerance potential of bacterial communities from soil and sediments along a gold tailing contamination gradient. Canadian journal of microbiology. 2017;63(9):788–805. https://doi.org/10.1139/cjm-2017-0214

31. Spain AM, Krumholz LR, Elshahed MS. Abundance, composition, diversity and novelty of soil Proteobacteria. The ISME journal. 2009;3(8):992–1000. https://doi.org/10.1038/ismej.2009.43

32. Sheik CS, Mitchell TW, Rizvi FZ, Rehman Y, Faisal M, Hasnain S, et al. Exposure of soil microbial communities to chromium and arsenic alters their diversity and structure. PloS one. 2012;7(6):e40059. https://dx.doi.org/10.1371%2Fjournal.pone.0040059

33. Huang C-C, Liang C-M, Yang T-I, Chen J-L, Wang W-K. Shift of bacterial communities in heavy metal-contaminated agricultural land during a remediation process. Plos one. 2021;16(7):e0255137. https://doi.org/10.1371/journal.pone.0255137

34. Riley TV, Mee BJ. Susceptibility of Bacteroides spp. to heavy metals. Antimicrobial agents and chemotherapy. 1982;22(5):889–92. https://doi.org/10.1128/aac.22.5.889

35. Li J, Mandal G, Rosen BP. Expression of arsenic resistance genes in the obligate anaerobe Bacteroides vulgatus ATCC 8482, a gut microbiome bacterium. Anaerobe. 2016;39:117–23. https://doi.org/10.1016/j.anaerobe.2016.03.012

36. Li P, Jiang D, Li B, Dai X, Wang Y, Jiang Z, et al. Comparative survey of bacterial and archaeal communities in high arsenic shallow aquifers using 454 pyrosequencing and traditional methods. Ecotoxicology. 2014;23(10):1878–89. https://doi.org/10.1007/s10646-014-1316-5

37. Wang Y, Li P, Jiang D, Li B, Dai X, Jiang Z, et al. Vertical distribution of bacterial communities in high arsenic sediments of Hetao Plain, Inner Mongolia. Ecotoxicology. 2014;23(10):1890–9. https://doi.org/10.1007/s10646-014-1322-7

38. Lv C, Zhao C, Yang S, Qu Y. Arsenic metabolism in purple nonsulfur bacteria. Wei sheng wu xue bao= Acta microbiologica Sinica. 2012;52(12):1497.

39. Wang J, Wu M, Lu G, Si Y. Biotransformation and biomethylation of arsenic by Shewanella oneidensis MR-1. Chemosphere. 2016;145:329–35. https://doi.org/10.1016/j.chemosphere.2015.11.107

40. Barton LL, Fardeau M-L, Fauque GD. Hydrogen sulfide: a toxic gas produced by dissimilatory sulfate and sulfur reduction and consumed by microbial oxidation. The Metal-Driven Biogeochemistry of Gaseous Compounds in the Environment: Springer; 2014. p. 237–77. https://doi.org/10.1007/978-94-017-9269-1_10

41. Weaver PF, Wall JD, Gest H. Characterization of Rhodopseudomonas capsulata. Arch Microbiol. 1975;105(1):207–16. https://doi.org/10.1007/bf00447139

42. Govarthanan M, Kamala-Kannan S, Selvankumar T, Mythili R, Srinivasan P, Kim H. Effect of blue light on growth and exopolysaccharides production in phototrophic Rhodobacter sp. BT18 isolated from brackish water. Int J Biol Macromol. 2019;131:74–80. https://doi.org/10.1016/j.ijbiomac.2019.03.049

43. Xiao E, Krumins V, Dong Y, Xiao T, Ning Z, Xiao Q, et al. Microbial diversity and community structure in an antimony-rich tailings dump. Applied microbiology and biotechnology. 2016;100(17):7751–63. https://doi.org/10.1007/s00253-016-7598-1

44. Lu H, Sato Y, Fujimura R, Nishizawa T, Kamijo T, Ohta H. Limnobacter litoralis sp. nov., a thiosulfate-oxidizing, heterotrophic bacterium isolated from a volcanic deposit, and emended description of the genus Limnobacter. International journal of systematic and evolutionary microbiology. 2011;61(2):404–7. https://doi.org/10.1099/ijs.0.020206-0

45. Spring S, Kämpfer P, Schleifer KH. Limnobacter thiooxidans gen. nov., sp. nov., a novel thiosulfate-oxidizing bacterium isolated from freshwater lake sediment. International journal of systematic and evolutionary microbiology. 2001;51(4):1463–70. https://doi.org/10.1099/00207713-51-4-1463

46. Zhang M, Wan K, Zeng J, Lin W, Ye C, Yu X. Co-selection and stability of bacterial antibiotic resistance by arsenic pollution accidents in source water. Environ Int. 2020;135:105351. Epub 2019/12/04. https://doi.org/10.1016/j.envint.2019.105351

47. Huaidong H, Waichin L, Riqing Y, Zhihong Y. Illumina-based analysis of bulk and rhizosphere soil bacterial communities in paddy fields under mixed heavy metal contamination. Pedosphere. 2017;27(3):569–78. http://dx.doi.org/10.1016/S1002-0160(17)60352-7

48. Bondici V, Lawrence J, Khan N, Hill J, Yergeau E, Wolfaardt G, et al. Microbial communities in low permeability, high pH uranium mine tailings: characterization and potential effects. Journal of applied microbiology. 2013;114(6):1671–86. https://doi.org/10.1111/jam.12180

49. Li J, Sun W, Wang S, Sun Z, Lin S, Peng X. Bacteria diversity, distribution and insight into their role in S and F e biogeochemical cycling during black shale weathering. Environ Microbiol. 2014;16(11):3533–47. https://doi.org/10.1111/1462-2920.12536

50. Wolfe-Simon F, Blum JS, Kulp TR, Gordon GW, Hoeft SE, Pett-Ridge J, et al. A bacterium that can grow by using arsenic instead of phosphorus. Science. 2011;332(6034):1163–6. https://doi.org/10.1126/science.1197258

51. Jain R, Jha S, Mahatma MK, Jha A, Kumar GN. Characterization of arsenite tolerant Halomonas sp. Alang-4, originated from heavy metal polluted shore of Gulf of Cambay. Journal of Environmental Science and Health, Part A. 2016;51(6):478–86. https://doi.org/10.1080/10934529.2015.1128717

52. Hamamura N, Itai T, Liu Y, Reysenbach AL, Damdinsuren N, Inskeep WP. Identification of anaerobic arsenite-oxidizing and arsenate-reducing bacteria associated with an alkaline saline lake in K hovsgol, M ongolia. Environmental Microbiology Reports. 2014;6(5):476–82. https://doi.org/10.1111/1758-2229.12144

53. Mukherjee P, Mitra A, Roy M. Halomonas rhizobacteria of Avicennia marina of Indian Sundarbans promote rice growth under saline and heavy metal stresses through exopolysaccharide production. Frontiers in microbiology. 2019;10:1207. https://doi.org/10.3389/fmicb.2019.01207

54. Katsoyiannis IA, Zouboulis AI. Application of biological processes for the removal of arsenic from groundwaters. Water research. 2004;38(1):17–26. https://doi.org/10.1016/j.watres.2003.09.011

55. Li P, Wang Y, Dai X, Zhang R, Jiang Z, Jiang D, et al. Microbial community in high arsenic shallow groundwater aquifers in Hetao Basin of Inner Mongolia, China. PloS one. 2015;10(5). https://doi.org/10.1371/journal.pone.0125844

